# Temperature-dependent estimation of Gibbs energies using an updated group contribution method

**DOI:** 10.1101/289918

**Authors:** Bin Du, Zhen Zhang, Sharon Gruber, James T. Yurkovich, Bernhard O. Palsson, Daniel C. Zielinski

## Abstract

Reaction equilibrium constants determine the mass action ratios necessary to drive flux through metabolic pathways. Group contribution methods offer a way to estimate reaction equilibrium constants at wide coverage across the metabolic network. Here, we present an updated group contribution method with: 1) additional curated thermodynamic data used in fitting; and 2) capabilities to calculate equilibrium constants as a function of temperature. We first collected and curated aqueous thermodynamic data, including reaction equilibrium constants, enthalpies of reaction, Gibbs free energies of formation, enthalpies of formation, entropies change of formation of compounds, and proton and metal ion binding constants. We further estimated magnesium binding constants for 618 compounds using a linear regression model validated against measured data. Next, we formulated the calculation of equilibrium constants as a function of temperature and calculated necessary parameters, including standard entropy change of formation (Δ_f_ *S*^∘^) and standard entropy change of reaction (Δ_r_*S*^∘^), using a model based on molecular properties. The median absolute errors in estimating Δ_f_ *S*^∘^ and Δ_r_*S*^∘^ were 0.010 kJ/K/mol and 0.018 kJ/K/mol, respectively. The efforts here fill in gaps for thermodynamic calculations under various conditions, specifically different temperatures and metal ion concentrations. These results support the study of thermodynamic driving forces underlying the metabolic function of organisms living under diverse conditions.

## Introduction

The First and Second Laws of Thermodynamics connect reaction flux directions, metabolite concentrations, and reaction equilibrium constants. An increasing number of systems biology methods have begun to take advantage of the intimate connection between thermodynamics and metabolism to obtain insights into the function of metabolic networks. These methods have been used in a number of applications including the calculation of thermodynamically-feasible optimal states (1, 2), the identification of thermodynamic bottlenecks in metabolism (3, 4), and the constraint of kinetic constants via Haldane relationships (5).

To perform thermodynamic analyses on metabolic networks, it is necessary to have values for the equilibrium constants of reactions carrying flux in the network. Experimentally, the equilibrium constant of a reaction is determined by calculating the mass action ratio (the ratio of product to substrate concentrations), also called the reaction quotient, when the reaction is at equilibrium. A collection of experimentally measured equilibrium constants for over 600 reactions has been published in the NIST Thermodynamics of Enzyme-Catalyzed Reactions database (TECRdb) (6). However, the equilibrium constants of the majority of known metabolic reactions are still missing, making computational estimation necessary. The most commonly used approach for estimating thermodynamic constants in aqueous solutions is the group contribution method (7, 8). This method is based on the simplifying assumptions that the Gibbs energy of formation (Δ_f_ *G*^∘^) of a compound is based on the sum of the contributions of its composing functional groups, which are independent of each other. The contribution of each group can be estimated through linear regression, using existing data on Δ_f_ *G*^∘^ and Gibbs energies of reactions (Δ_r_*G*^∘^).

Recent iterations of group contribution methods for reactions in aqueous solutions have incorporated pH corrections into estimations of equilibrium constants (9) and improved accuracy by taking advantage of fully-defined reaction stoichiometric loops forming First Law energy conservation relationships within the training data (10). These methods also have begun to take advantage of computational chemistry software to estimate the p*K*_a_s of compounds as part of thermodynamic parameter estimation. However, a number of issues remain for thermodynamic estimation of reaction equilibrium constants in metabolic networks, including: 1) significant estimation errors in many cases, which may be attributed to a number of factors including missing or erroneous reaction conditions, and 2) the lack of an established method to handle correction of thermodynamic data with respect to temperature changes across conditions. Additionally, existing group contribution methods have not taken into account the substantial metal ion binding of many metabolites at physiological ion concentrations, although established theory exists to correct reaction equilibrium constants for metal ion binding when ion dissociation constants are available (11).

In this study, we extend the capabilities of computational estimation of reaction equilibrium constants for metabolic networks. We first curate the NIST TECRdb of reaction equilibrium constants to obtain missing reaction conditions and correct any other errors. We further incorporate additional thermodynamic data, including data related to proton and metal ion binding, from a number of other sources (12–14). To fill gaps in metal binding correction of equilibrium constants, we estimate magnesium binding constants for 618 compounds using molecular descriptors and magnesium binding groups defined based on known magnesium binding compounds. We also optimize magnesium binding constants using equilibrium constants measured at different magnesium concentrations through a least-squares method. Next, to enable calculation of equilibrium constants as a function of temperature, we adapt the thermodynamic theory from the geochemistry literature (15–21) given certain simplifying assumptions. The thermodynamic parameters required for such calculation, Δ_f_ *S*^∘^ and Δ_r_*S*^∘^, are estimated through regression models using various molecular descriptors. Finally, we incorporate these new data and functionalities into the most recently published group contribution framework, termed the component contribution (10), to obtain a new group contribution estimator for reaction equilibrium constants with expanded capabilities.

## Materials and Methods

### Workflow for estimation of equilibrium constants

A total of 4298 equilibrium constants (*K*′) for 617 unique reactions measured under different conditions (temperature, pH, ionic strength, metal ion concentrations) were used as the training data set for the current group contribution method. First of all, we transformed all measurements to the same reference conditions at 298.15 K, pH 7, 0 M ionic strength and no metal concentration. We applied a Legendre transform to account for the different ion binding states of each compound as in the previous component contribution method (10). The transformation of Gibbs free energy of reaction across pH and ionic strength is also based on the previous method, except that we used the Davies equation rather than Debye-Hückel equation to calculate activity coefficients of electrolyte solutions. The transformation of Gibbs free energy of reaction across different metal concentrations is based on the formulation described by Alberty (11, 22). The transformation of Gibbs free energy of reaction across temperature is based on adapted thermodynamic theory from the geochemistry literature (15–18) with simplifying assumptions. The relevant equations and theory above can be found in the Supporting Material.

Using Δ_r_*G*^∘^ and Δ_f_ *G*^∘^ data at reference conditions, we applied the component contribution method by Noor et al (10) and obtained estimates of Δ_r_*G*^∘^ and Δ_f_ *G*^∘^ at reference conditions. Using these values, we are able to calculate the equilibrium constant of a given reaction at defined conditions by applying the transformations mentioned earlier.

### Curation of The IUPAC Stability Constants Database

The IUPAC Stability Constants Database (SC-database) contains ion binding data, i.e. dissociation/binding/stability constants, under various conditions from primary literature. Additionally, the database contains several different annotations for binding of protons and metal ions to specific aqueous species. When the ligand is a proton, the related dissociation constant is a p*K*_a_ constant, while when the ligand is a metal ion such as magnesium, the dissociation constant is a p*K*_Mg_ (modified to the specific ion) constant. For each compound of interest, we categorized the available binding data specific to each ion bound state. We then corrected binding data to 0 M ionic strength using the Davies equation (23). For each ion binding reaction, we calculated the median of all available binding data as the value utilized in the fitting (Table S4, S5 and S7).

### Features and data used in regression models

For estimation of magnesium binding constants (p*K*_Mg_), we included a total of 140 data points (Table S5) and 128 molecular descriptors as features for regression models. The molecular descriptors included metal binding groups identified from existing p*K*_Mg_ data (Table S8), the charge of the compound excluding any metal binding groups, sums of partial charge and numbers of different types of atoms, and several additional molecular descriptors from ChemAxon and RDkit. For estimation of Δ_f_ *S*^∘^, we included 669 data points (Table S3) and 186 features including group decompositions, sums of partial charge and numbers of different types of atoms, and molecular descriptors from ChemAxon and RDkit. For estimation of Δ_r_*S*^∘^, we included 617 data points (Table S10) and 223 features including group decompositions, Δ_r_*G*^∘^ of the reaction at 298.15 K and difference in product and substrate number, sums of partial charge and numbers of different types of atoms, and molecular descriptors from ChemAxon and RDkit. The molecular descriptors of compound were estimated with Calculator Plugins, Marvin 16.11.21, 2016, ChemAxon (http://www.chemaxon.com) and RDKit: Open-source cheminformatics (http://www.rdkit.org). The full list of molecular descriptors that were used can be found in Table S16.

### Comparison of regression methods using nested 10-fold cross-validation

We applied nested 10-fold cross-validation to compare the performance of various regression methods for estimation of p*K*_Mg_, Δ_f_ *S*^∘^ and Δ_r_*S*^∘^. The methods tested were ridge regression, lasso regression, elastic net regularization, random forests, extra trees and gradient boosting. The specific implementation of nested 10-fold cross-validation involves generating an outer loop and inner loop of cross-validation. The outer loop separates the whole dataset into 10 folds, with one fold for testing and the rest for training in each iteration. The training data in each iteration is further separated into 10-folds, and cross-validation is performed in the inner loop to select the optimal hyperparameters through grid search (Table S17). We repeated the nested 10-fold cross-validation on each regression method 5 different times by splitting the data into different subdivisions.

We then assessed model performance through the median absolute residual of testing errors calculated from the outer loop, for a total of 50 folds (10 folds × 5 repetitions). The testing errors calculated here also reflect how well the model generalizes on unseen data and are thus used as a metric to evaluate model performance. We also evaluated model stability by calculating the relative standard deviation (RSD = standard deviation/mean) of hyperparameters selected by the inner loop, for a total of 50 folds (10 folds × 5 repetitions). We selected the regression model that has the lowest testing error and RSD of hyperparameters. For every fitting procedure, we applied standardization on both the training and testing set using the mean and standard deviation of features calculated from the training set.

The regression models, including linear models and tree-based methods, were implemented using the python package scikit-learn 0.19.1 (24).

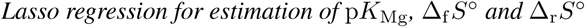

We used lasso regression to estimate p*K*_Mg_, Δ_f_ *S*^∘^ and Δ_r_*S*^∘^. Specifically, the objective function to minimize is

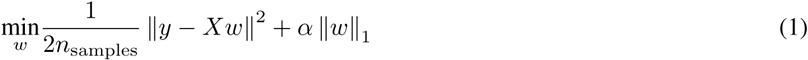

where *y* is the vector of data with length *n*_samples_, *X* is the matrix with features in the row corresponding to each data point, *w* is the vector of coefficients of the model, *α* is a constant that tunes the degree of *l*_1_ penalty.

We repeated 10-fold cross-validation 100 times on p*K*_Mg_, Δ_f_ *S*^∘^ and Δ_r_*S*^∘^ datasets respectively to find the optimal *α* values that lead to the lowest testing errors. We then constructed a lasso regression-based estimator for each p*K*_Mg_, Δ_f_ *S*^∘^ and Δ_r_*S*^∘^ dataset, using the selected *α* value and applying standardization on the dataset.

### Optimization of ion binding constants using the Levenberg-Marquardt algorithm

For selected reactions with compound ion binding constants to optimize, we first collected *K*× data measured at different magnesium concentrations. We then formulated equations that correct the standard transformed Gibbs energy of reaction (Δ_r_*G*′^∘^) (calculated from *K*×) to Δ_r_*G*^∘^, where magnesium binding constants are variables in the equations. We allowed ±0.5 variation for each ion binding constant from its original value, consistent with reported error in these parameters (25, 26). We optimized the binding constants using an iterative Levenberg-Marquardt algorithm with decreasing step sizes for the gradient approximation parameter. At each iteration, we input the optimized values from the previous iteration into the transformation equations and calculated the least-squares errors in estimating Δ_r_*G*^∘^. The termination criterion for the optimization was a fractional difference in the sum of squares between two consecutive iterations below 0.00001 (unitless). The Levenberg-Marquardt algorithm was performed using python package lmfit 0.9.2 (27).

### Comparison of previous and current group contribution method

We used 10-fold cross-validation on the 432 reactions that overlapped between the previous (10) and the current group contribution method. Specifically, we first transformed experimentally measured Δ_r_*G*^*/*∘^ data to the reference state Δ_r_*G*^∘^ (298.15 K, pH 7, 0 M ionic strength), calculated the median Δ_r_*G*^∘^ of all data points in each unique reaction, and performed 10-fold cross-validation on those 432 Δ_r_*G*^∘^ values. We repeated this procedure 100 times by splitting the data into different subdivisions. We then calculated the median absolute residual of 100 repetitions for each reaction.

Additionally, we also compared how well the two methods perform on the 185 new reactions collected in this work. Specifically, we fit the group contribution model using both methods with Δ_r_*G*^∘^ values of the original 432 overlapping reactions as training data, and calculated the absolute residual in predicting Δ_r_*G*^∘^ for the 185 new reactions as the testing set.

### Calculation of standard entropy change of formation

The standard entropy change of formation (Δ_f_ *S*^∘^) of the compound is not directly available. It can be calculated either from Δ_f_ *G*^∘^ and and the standard enthalpy of formation (Δ_f_ *H*^∘^) of the compound

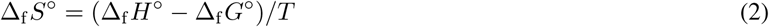

or from the standard molar entropy (*S*^∘^) of the compound

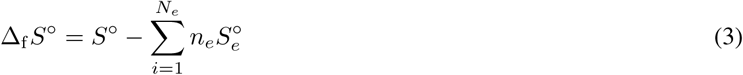

where 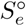 is the standard molar entropy of the element *N*_*e*_ composing the compound and *n*_*e*_ is the number of atoms for the element *N*_*e*_.

### Implementation and availability of source code

The updated group contribution method has been implemented in python 2.7.6. The source code is available on GitHub (https://github.com/bdu91/group-contribution), together with detailed instructions on how to install and examples using the package.

## Results

### Collection and curation of thermodynamic data

The workflow for estimating reaction equilibrium constants under given pH, temperature, ionic strength and metal ion concentrations is demonstrated in Figure 1A (Materials and Methods). To obtain the necessary data for this estimation, we curated a number of databases and primary literature sources. First of all, from NIST TECRdb (https://randr.nist.gov/enzyme) (6), we obtained measured equilibrium constants (*K*′) and enthalpies of reactions (Δ_r_*H*′^∘^) for 617 and 207 unique reactions, respectively. Noticing a number of gaps in experimental conditions and other minor issues, we curated a total of 4298 measured *K*′ data from NIST TECRdb. This curation effort resulted in 48.9% corrected data entries, including updated experimental media conditions (35.78%), addition of new data (5.12%), correction of *K*′ values (3.49%), removal of problematic data (3.33%) (examples in Table S14) and correction of reaction formulae (1.14%) (Figure 1B).

**Figure 1:**
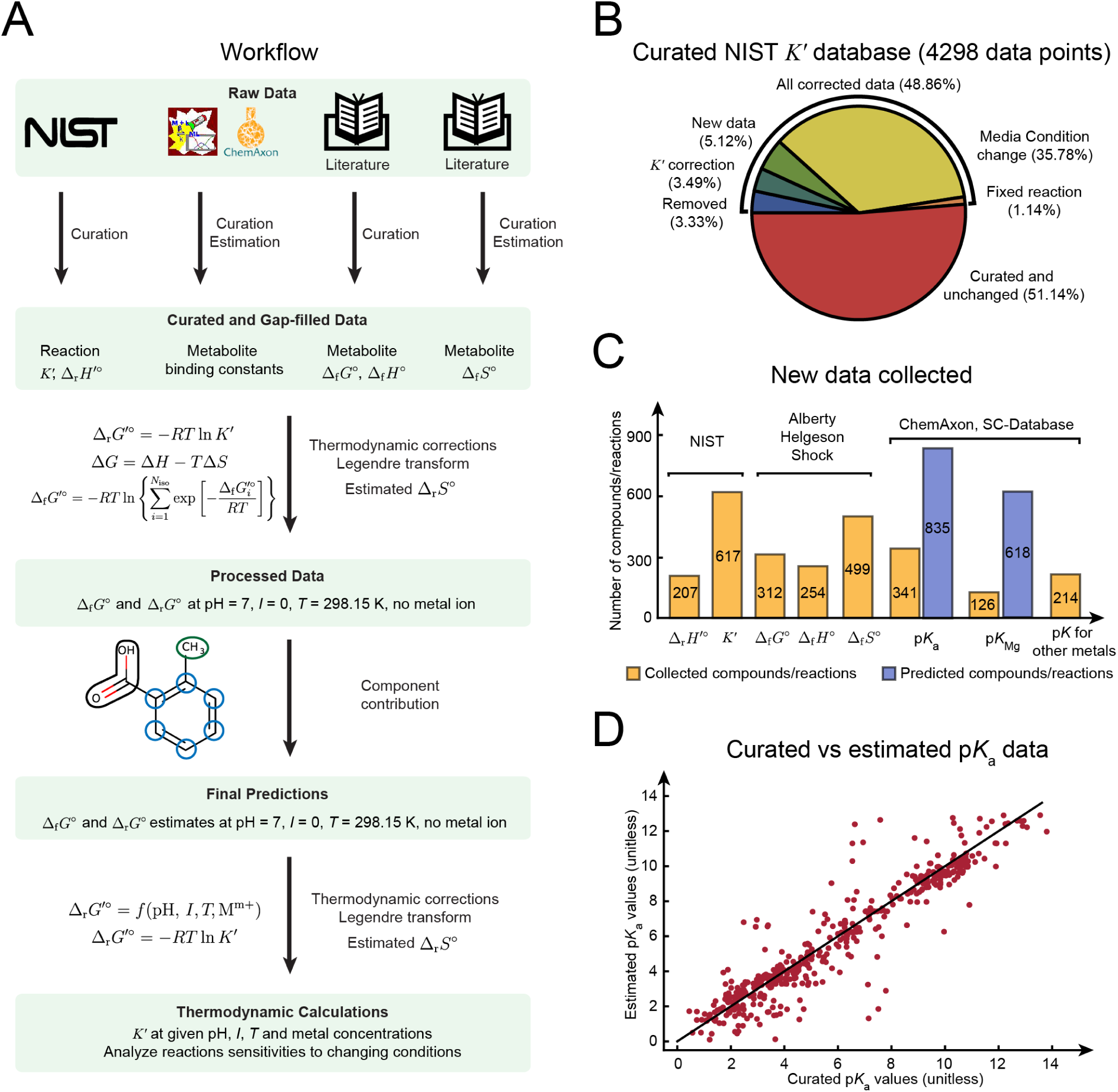
Estimation of reaction equilibrium constants. (A) Workflow of data curation and parameter fitting for equilibrium constant estimation. (B) The results of curation of equilibrium constants from the NIST Thermodynamics of enzyme-catalyzed reactions database (TECRdb). (C) New thermodynamic properties generated, either collected from sources shown or computationally estimated. (D) Comparison between curated p*K*_a_ data from The IUPAC Stability Constants Database (SC-database) as well as literature with computationally estimated p*K*_a_ values from ChemAxon.

Next, we collected data on standard Gibbs free energies of formation (Δ_f_ *G*^∘^), standard enthalpies of formation (Δ_f_ *H*^∘^) and standard entropy of formation changes (Δ_f_ *S*^∘^) for 312, 254 and 499 unique compounds, respectively (Figure 1C). The above data are from multiple sources: *Thermodynamics of Biochemical Reactions* by Alberty (11), the SUPCRT92 database (28) and Organic Compounds Hydration Properties Database (29). It is worth mentioning that Δ_f_ *S*^∘^ data are usually not directly measured but instead are calculated from either Δ_f_ *G*^∘^ and Δ_f_ *H*^∘^ data or standard molar entropy (*S*^∘^) of the compound (Materials and Methods).

Lastly, we collected and curated p*K*_a_ data for 341 compounds, magnesium binding constants for 126 compounds, and other metal type binding constants for 214 compounds (including cobalt, iron, zinc, sodium, potassium, manganese, calcium, lithium) from The IUPAC Stability Constants Database (SC-database) and a few primary literature (12–14) (Figure 1C). We also predicted p*K*_a_ data for 835 compounds using ChemAxon (http://www.chemaxon.com) (Figure 1C). Differences between collected p*K*_a_ data and the predicted values from ChemAxon for the same compounds can be as large as 5.84 (unitless), with a median of 0.42 (unitless) (Figure 1D). This error is a large enough difference to substantially alter the major protonation states for metabolites containing groups with p*K*_a_s around physiological pH, such as phosphate groups. We examined the specific cause of the largest discrepancies and found that they are due to issues such as assignment of p*K*_a_ value to the wrong charged form by ChemAxon (e.g. 4-oxo-L-proline) or error in calculating p*K*_a_s related to particular molecular moieties, such as nitrogenous bases and nitrogen atoms on unsaturated rings (e.g. 2’-deoxyguanosine 5’-monophosphate, xanthine-8-carboxylate, deaminocozymase). We thus used measured p*K*_a_ data when available. All collected and curated data can be found in the Supporting Material (Table S1-7).

### Estimation of magnesium binding constants

In aqueous solutions, the standard transformed Gibbs free energy of the compound (Δ_f_ *G*′^∘^) can depend on pH and concentrations of metal ions, due to the presence of different protonation states and various metal bound species. Specifically, Δ_f_ *G*′^∘^ can be calculated based on the standard transformed Gibbs energies of its different ion bound states (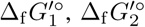, etc) through Legendre transform (11).

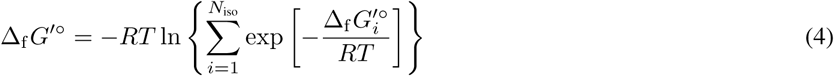

The equation can be rewritten as

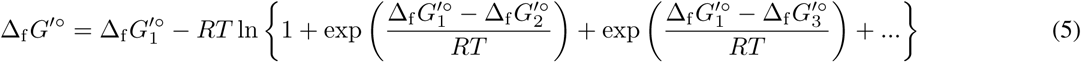

where 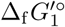 is the Gibbs energy of a particular ion bound state (typically with the least hydrogens and metal ions bound). Thus, the Gibbs energy of a specific ion bound state 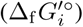 can be written in terms of 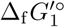 and the binding polynomial *P*_*i*_

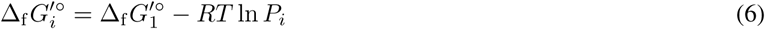

where *P*_*i*_ is expressed in terms of the proton concentration and metal ion concentration, as well as the binding constants of successive proton and metal ion binding steps to obtain the *i*^*th*^ ion bound state (11) (derivation in Supporting Material). Therefore, metal binding constants are important parameters that affect Δ_f_ *G*′^∘^ and subsequently reaction equilibrium constants.

We focused on magnesium binding since magnesium ion is well known to bind to various metabolites, and its binding to ATP and other phosphate-containing compounds has been characterized experimentally (30, 31). However, magnesium binding data is still lacking for a large number of compounds that contain similar structural groups to those known to bind magnesium, suggesting that many more compounds may have substantial magnesium binding than have been measured.

Based on the structures of compounds with known magnesium binding, we determined 31 magnesium binding groups (Table S8), most of which are phosphate and carboxyl groups. Together with other molecular descriptors (Materials and Methods), we used 128 features and 140 measured magnesium binding constants to construct several candidate regression models for the prediction of magnesium dissociation constants. We performed nested 10-fold cross-validation to compare between multiple regression models (Figure S2A-B). We selected lasso regression as the best predictor due to its superior generalizability compared to more complex methods (Figure S2A) and stable model parameters across cross-validation replicates compared to other linear methods (Figure S2B). Using 140 measured magnesium binding constants as training data, we constructed a lasso regression model with parameters tuned through cross-validation (Figure 2A) and predicted 1707 magnesium binding constants for aqueous species from 618 compounds (Table S5). The most predictive variables in the model include the formal charge, the solvent accessible surface area, the presence of various phosphate groups for magnesium binding, the partial charge of nitrogen atoms, the compound charge excluding its magnesium binding groups and dipole moment of the molecule. We found 34 of the 618 compounds are predicted to bind to magnesium at physiological concentrations (2 to 3 mM) (32). The median absolute residual of the lasso regression model for magnesium binding constant estimation is 0.39 (unitless), as calculated by the nested 10-fold cross-validation (Figure S2A).

**Figure 2:**
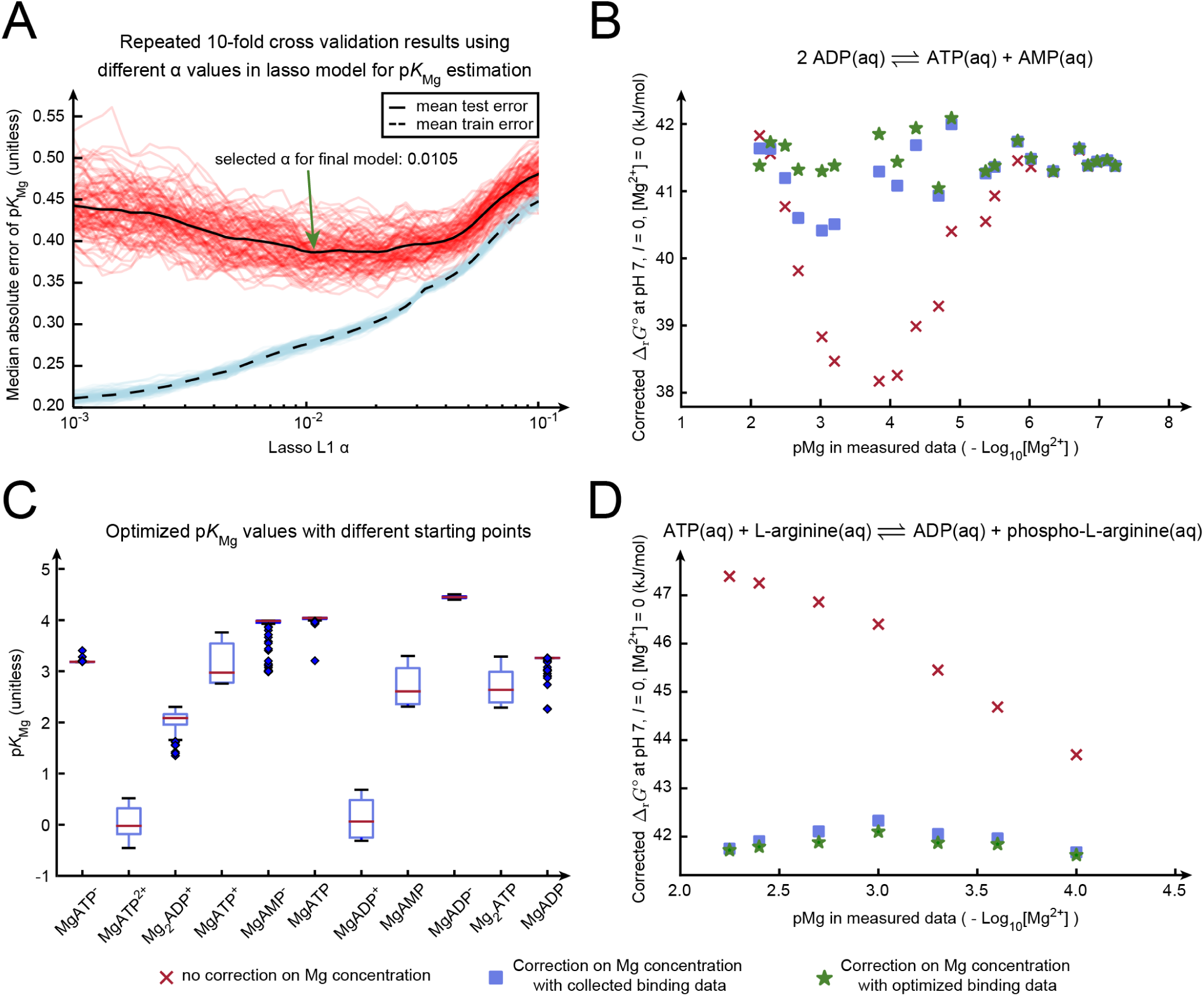
Estimation of compound magnesium binding constants (p*K*_Mg_). (A) Selection of parameters in the lasso regression using 10-fold cross-validation on all p*K*_Mg_ data. We repeated 10-fold cross-validation 100 times and calculated training (blue) and testing (red) errors at *α* from 10^−3^ to 10^−1^. The mean training and testing errors are shown in dashed and solid black lines. The selected *α* at the lowest mean testing error is 0.0105 (unitless). (B) Corrected Δ_r_*G*^∘^ values of the adenylate kinase reaction calculated from equilibrium constants (*K*′) measured at different Mg concentrations. We applied different corrections to transform Δ_r_*G*′^∘^ (calculated from *K*′) to Δ_r_*G*^∘^ values: no correction on varying Mg concentrations (red Xs), a correction on Mg concentrations using collected p*K*_Mg_ data (blue squares), and a correction on Mg concentrations using p*K*_Mg_ values optimized through a least-squares method (green stars). (C) Optimized p*K*_Mg_ values from adenylate kinase reaction with different starting points. We randomly selected p*K*_Mg_ within ±0.5 of its original value as the starting point for the least-squares optimization. We repeated this procedure 100 times and showed the range of optimized p*K*_Mg_ values from those iterations. (D) Corrected Δ_r_*G*^∘^ values of the arginine kinase reaction from equilibrium constants (*K*′) measured at different Mg concentrations. The markers are the same as in panel B, except that green stars represent corrected Δ_r_*G*^∘^ values using optimized p*K*_Mg_ from adenylate kinase reaction data.

### *Optimization of magnesium binding constants using* Δ_r_*G*′^∘^ *data*

To assess the quality of magnesium binding data, we examined how well the binding constants correct *K*′data measured at different magnesium concentrations to the same reference conditions. Specifically, we transformed the Δ_r_*G*′^∘^ data calculated from *K*′ (Δ_r_*G*′^∘^ = *RT* ln *K*′) to the reference state Δ_r_*G*^∘^ (298.15 K, pH 7, 0 M ionic strength, no metal ion) using Legendre transforms (Supporting Material) (11). The resulting reference state Δ_r_*G*^∘^ values should be within a small range, which would indicate that the correction to Gibbs energy for magnesium binding is accurate. Taking data from the reaction catalyzed by adenylate kinase, one of the best characterized reactions, as an example, we found a substantial decrease in Δ_r_*G*^∘^ variation with respect to magnesium concentration after applying corrections to account for magnesium binding (Figure 2B). However, in some cases, the differences in Δ_r_*G*^∘^ remained substantial. This issue indicates that significant error may exist in certain measured binding constants, likely due to a combination of experimental complications during measurements as well as the fact that several different binding constants often exist for the same compound, making it difficult to infer individual binding constants from experimental data with high confidence.

To address the errors associated with measured binding constants, we attempted to adjust the binding constants to maximize the correction on *K*′ data at different magnesium concentrations. Specifically, we optimized the binding constants of adenylate kinase reaction using Levenberg-Marquardt algorithm that minimizes the least-square errors on Δ_r_*G*^∘^ (33, 34). When we applied the optimized binding constants (Table S6) to the data for adenylate kinase reaction, we found Δ_r_*G*^∘^ values now fell within a smaller range (Figure 2B) compared to Δ_r_*G*^∘^ values calculated from pre-optimized binding constants, indicating improved performance of the optimized binding constants. To assess global optimality of the solution, we randomly selected magnesium binding constants within ±0.5 of their original values as starting points and repeated the algorithm 100 times. We found that optimized binding constants are generally stable, although a number of them have some degree of uncertainty (Figure 2C). On the other hand, Δ_r_*G*^∘^ values calculated using those different sets of optimized binding constants did fall in similar ranges (Figure 2B).

To test how these optimized binding constants work on other datasets, we applied them on Δ_r_*G*′^∘^ data from arginine kinase reaction (Figure 2D), and obtained a decrease of variation of Δ_r_*G*^∘^ values with respect to magnesium concentration compared to those calculated from pre-optimized binding constants. We observed similar trend when applying magnesium binding constants optimized from citrate hydrolase reaction data to Δ_r_*G*′^∘^ data from aconitase reaction (Figure S3D, S3E). In total we examined nine reactions in NIST TECRdb that exhibited clear trends in *K*′ with respect to magnesium ion concentration (Figure 2, S2). In the majority of these cases, applying the collected/estimated magnesium binding constants reduces variation in Δ_r_*G*^∘^ (Figure S2 yellow background). In cases where the estimated binding constants did not reduce variation in Δ_r_*G*^∘^ (Figure S2 grey background), we applied the Levenberg-Marquardt algorithm to minimize the least-square errors on Δ_r_*G*^∘^ to obtain optimized binding constants that better correct the data. The optimized binding constants from all case studies can be found in Table S6, and statistics on case studies can be found in Table S12.

### *Thermodynamic parameters for transformation of* Δ_r_*G*′^∘^ *across temperature*

We then sought to develop the capability to calculate standard transformed Gibbs energy of reaction (Δ_r_*G*′^∘^) as a function of temperature. Specifically, we adapted theory from the geochemistry literature under constant enthalpy and entropy assumptions (15–18) to obtain a simple linear formulation of Δ_r_*G*′^∘^ at a given temperature *T* using the standard entropy change of reaction Δ_r_*S*^∘^ at a reference *T*_*r*_ (298.15 K) (derivation provided in the Supporting Material):

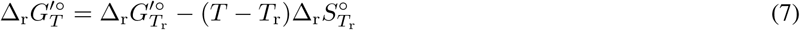

To estimate 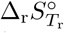 (we use Δ_r_*S*^∘^ in later references since *T*_*r*_ is the only condition of interest) of reactions, we first collected Δ_r_*S*^∘^ from multiple sources as training data. From NIST TECRdb, we selected reactions with *K*^*/*^ data measured under at least 4 different temperatures. We then calculated Δ_r_*S*^∘^ of each reaction using the Δ_r_*G*^*/*∘^ of the reaction at different temperatures based on Equation 7, obtaining 51 Δ_r_*S*^∘^ values. Next, we picked reactions in NIST TECRdb with both Δ_r_*G*^∘^ and Δ_r_*H*^∘^ data available and calculated their Δ_r_*S*^∘^ values, obtaining 41 additional data points. We also collected 669 Δ_f_ *S*^∘^ values for 499 compounds at different protonation states (Table S3) and obtained 56 Δ_r_*S*^∘^ values using Δ_f_ *S*^∘^ data. We noted that these 148 reactions with Δ_r_*S*^∘^ data available (Table S10) remain a small set compared to the 617 reactions in NIST TECRdb and cover only 61 groups out of a total of 106 groups. To obtain more training data for Δ_r_*S*^∘^ estimation, we sought to estimate Δ_f_ *S*^∘^ that can be used for Δ_r_*S*^∘^ calculation. The workflow for this estimation is divided into two parts: 1) estimation of Δ_f_ *S*^∘^ using collected 669 Δ_f_ *S*^∘^ values through regression model, 2) assessment of the regression model performance with the addition of Δ_r_*S*^∘^ values calculated from estimated Δ_f_ *S*^∘^.

### *Estimation of standard entropy change of formation* Δ_f_ *S*^∘^

We found that simple molecular descriptors, notably the number of atoms in the compound and the compound charge, were highly useful as predictors for Δ_f_ *S*^∘^. Specifically, we found Δ_f_ *S*^∘^ data to be highly correlated simply with the total numbers of atoms in the compound, with an *R*^2^ of 0.89 (Figure 3A). The Δ_f_ *S*^∘^ data as a function of atom number are separated into two main clusters, one of which contains aqueous species with large atom numbers and large absolute Δ_f_ *S*^∘^ values (NAD, NADH, NADP, NADPH). The other cluster contains a wide variety of aqueous species, with a few categories labeled in Figure 3A. We noticed clear separations among aqueous species with -5, -4, -3 and -2 charge, but less so for those with -1, 0 and +1 charge (Figure 3A). We found the trend between Δ_f_ *S*^∘^ and number of atoms exists even more strongly among compounds within the same homologous series, where the compound structures differ only by the number of CH_2_ units in the main carbon chain. Specifically, Δ_f_ *S*^∘^ value decreases by 0.11 kJ/K/mol with every additional CH_2_ unit. This trend was observed in a number of homologous series including alkanes, alkenes, alkynes, aldehydes, single carboxylic acids, amines, amides, and thiols. However, the change in Δ_f_ *S*^∘^ with respect to number of atoms across different homologous series is inconsistent, thus requiring additional molecular descriptors.

**Figure 3:**
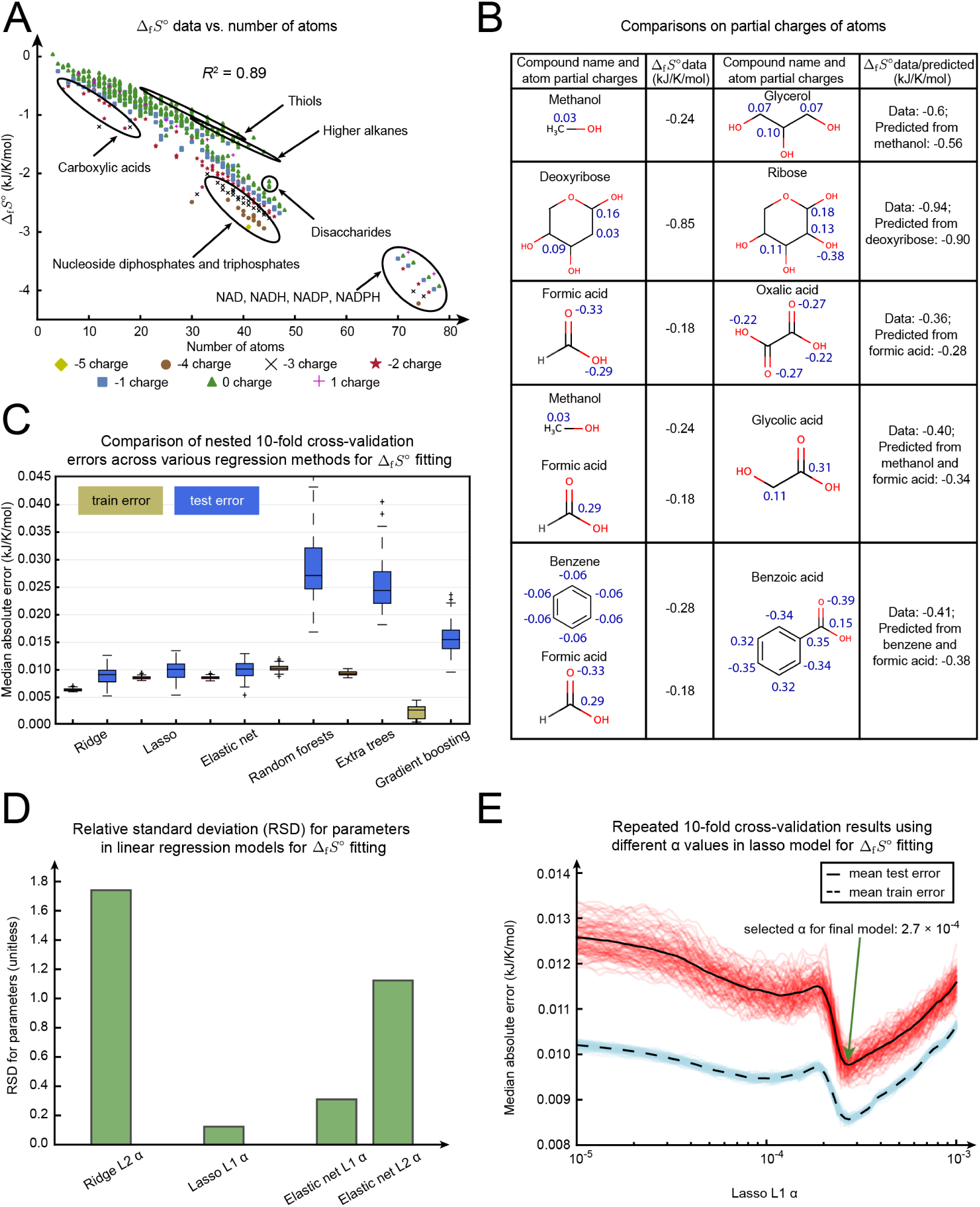
Estimation of standard entropy change of formation (Δ_f_ *S*^∘^). (A) Δ_f_ *S*^∘^ vs. number of atoms. The compounds with different charges are encoded with different colored symbols. We also labeled compounds belonging to the same category or containing the same functional groups. (B) Comparisons of partial charges of atoms between compounds. Each row contains a pair of compounds and their Δ_f_ *S*^∘^ data. In each pair, Δ_f_ *S*^∘^ of the latter compound can be predicted from that of the former compound(s) based on difference in atom number. The partial charges of atoms that are different within each pair are marked in blue. (C) Training and testing errors of nested 10-fold cross-validation (repeated 5 times) on Δ_f_ *S*^∘^ data using 6 different regression methods. (D) Relative standard deviation (RSD) of parameters using linear regression models for Δ_f_ *S*^∘^ fitting. We used RSD (standard deviation/mean) to assess the relative variability of parameters selected by the inner loops of nested cross-validation (repeated 5 different times). (E) Selection of parameters in the lasso regression using 10-fold cross-validation on all Δ_f_ *S*^∘^ data. We repeated 10-fold cross-validation 100 times and calculated training (blue) and testing (red) errors at *α* from 10^−5^ to 10^−3^. The mean training and testing errors are shown in dashed and solid black lines. The selected *α* at the lowest mean testing error is 2.7 × 10^−4^ (unitless).

As an additional descriptor, we found that partial charge of atoms can help distinguish Δ_f_ *S*^∘^ from different homologous series. For example, the carbon atoms in glycerol (alcohol containing multiple hydroxyl groups) have larger partial charges than those in methanol (alcohol containing a single group). The prediction of glycerol Δ_f_ *S*^∘^ from methanol Δ_f_ *S*^∘^ based on their difference in atom numbers yielded a smaller absolute Δ_f_ *S*^∘^ value than the actual glycerol data (Figure 3B) (calculation in Table S9). The correlation of larger partial charges of carbon atoms with larger absolute Δ_f_ *S*^∘^ is also observed in other pairs in Figure 3B (deoxyribose vs. ribose, methanol + formic acid vs. glycolic acid, benzene + formic acid vs. benzoic acid). Besides carbon atoms, we also found differences in partial charges of oxygen atoms to be associated with Δ_f_ *S*^∘^ differences, as shown between formic acid and oxalic acid (Figure 3B). Following these observations, we included the sums of absolute partial charge of each type of atom as molecular descriptors for the regression model.

In addition to partial charge, we also considered a number of other molecular descriptors from ChemAxon and RDkit (Materials and methods). We obtained a total of 186 features and 669 Δ_f_ *S*^∘^ data for regression models. We performed nested 10-fold cross-validation to compare between multiple regression models (Figure 3C, 3D). We selected lasso regression as the final model to use since it has significantly smaller testing errors compared to more complex methods (Figure 3C) and the least variation in parameters selected from cross-validation compared to other linear regression methods (Figure 4D). Using parameters selected from cross-validation on the entire Δ_f_ *S*^∘^ dataset (Figure 3E), we constructed a lasso regression model and predicted 672 Δ_f_ *S*^∘^ values (Table S3). The most predictive variables in the model included the number of hydrogen, nitrogen, phosphate and oxygen atoms, the partial charge of oxygen atoms, the formal charge of the compound, the presence of phosphate groups, the intramolecular steric hindrance, and the solvent accessible surface area. The median absolute residual of the lasso regression model for Δ_f_ *S*^∘^ estimation is 0.010 kJ/K/mol (Figure 3C).

**Figure 4:**
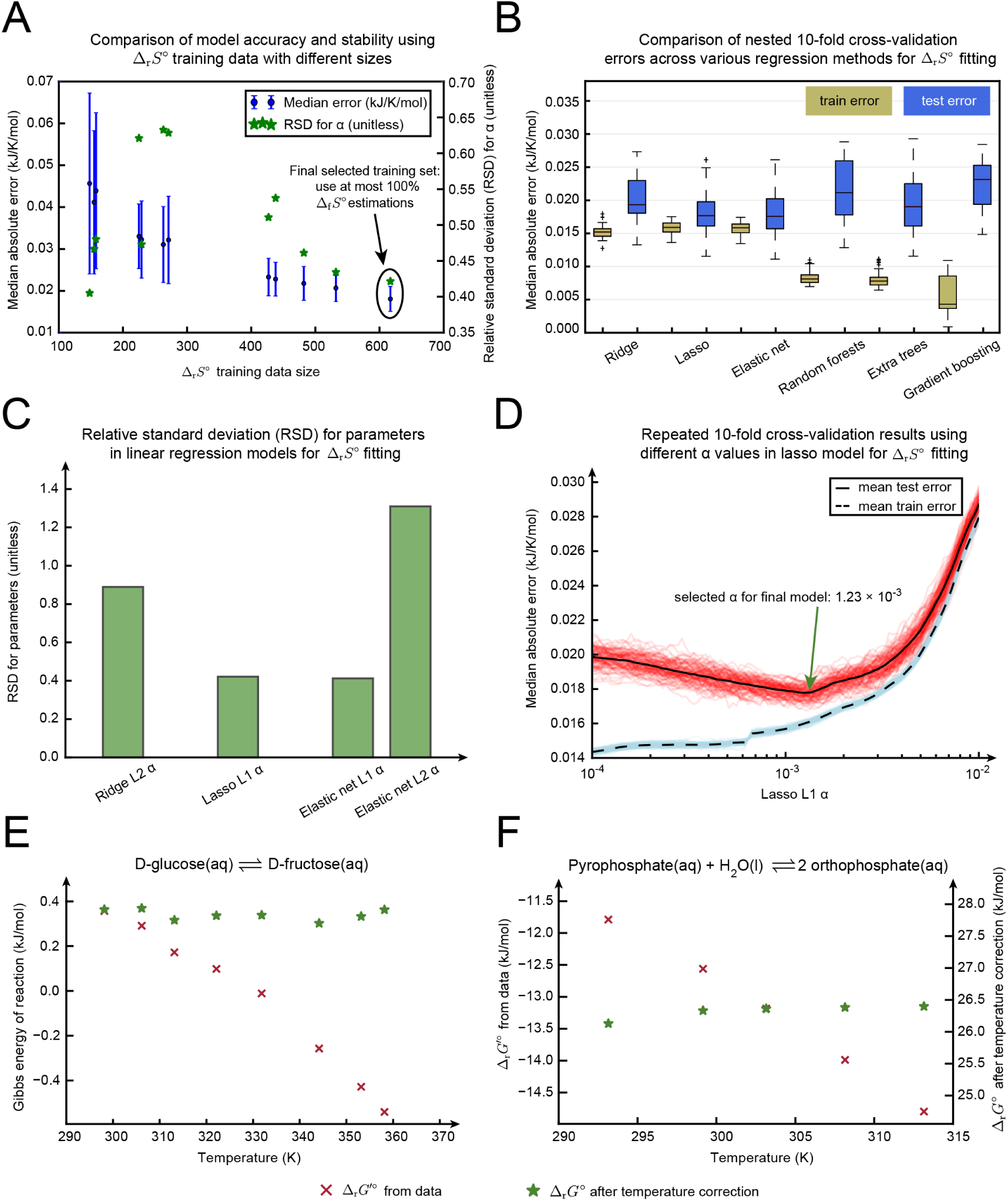
Estimation of standard entropy change of reaction (Δ_r_*S*^∘^). (A) Nested 10-fold cross-validation using the lasso regression on Δ_r_*S*^∘^ training data with different sizes. We obtained different Δ_r_*S*^∘^ training data sizes by setting cutoff at different fractions of estimated Δ_f_ *S*^∘^ used to calculate Δ_r_*S*^∘^. We performed this cross-validation five times and calculated the testing error (left y axis) and relative standard deviation (RSD) of *α* selected by inner loops of nested cross-validation (right y axis) at each cutoff. (B) Training and testing errors of nested 10-fold cross-validation (repeated 5 times) on 617 Δ_r_*S*^∘^ data using 6 different regression methods. (C) RSD of parameters selected by inner loops of nested cross-validation using linear regression models for Δ_r_*S*^∘^ fitting. (D) Selection of parameters in the lasso regression using 10-fold cross-validation on all 617 Δ_r_*S*^∘^ data. We repeated 10-fold cross-validation 100 times and calculated training (blue) and testing (red) errors at *α* from 10^−4^ to 10^−2^. The selected *α* at the lowest mean testing error is 1.23 × 10^−3^ (unitless). (E) Corrected Δ_r_*G*^∘^ (green stars) vs. Δ_r_*G*′^∘^ (red Xs) values (calculated from *K*^*/*^) of glucose isomerase reaction measured at different temperatures. (F) Corrected Δ_r_*G*^∘^ (green stars) vs. Δ_r_*G*′^∘^ (red Xs) values of pyrophosphatase reaction measured at different temperatures.

### *Estimation of standard entropy changes of reactions* Δ_r_*S*^∘^

In addition to the 148 Δ_r_*S*^∘^ training data points mentioned in earlier section, we expanded the training data by including Δ_r_*S*^∘^ calculated using both Δ_f_ *S*^∘^ data and estimation. Considering all possible fractions of Δ_f_ *S*^∘^ estimations used in a single reaction (0, 17%, 20%, 25%, 29%, 34%, 40%, 50%, 60%, 67%, 75%, 100%), we constructed multiple lasso regression models with different sizes of Δ_r_*S*^∘^ training data. We used 223 molecular descriptors as features for all regression models (Materials and Methods). Using nested 10-fold cross-validation, we found that the model with the largest number (617) of training data points (thus including Δ_r_*S*^∘^ calculated from all Δ_f_ *S*^∘^ estimations) has the smallest error and most stable parameters (RSD for *α*) selected from cross-validation (Figure 4A). To confirm the performance of lasso regression model for Δ_r_*S*^∘^ estimation, we performed nested 10-fold cross-validation on multiple regression models using 617 Δ_r_*S*^∘^ training data (Figure 4B, 4C). We found lasso regression has the lowest testing error (Figure 4B) and resulted in the most stable parameters (Figure 4C). Finally, with parameters selected from cross-validation on 617 Δ_r_*S*^∘^ training data (Figure 4D), we constructed a lasso regression model for Δ_r_*S*^∘^ estimation. The most predictive variables in the model included the number of atoms, the number of bonds, the formal charge of the compound, the partial charge of phosphate and oxygen atoms, the difference in the product and substrate number, the presence of phosphate groups and dipole moment of the molecule. The median absolute residual of the lasso regression model for Δ_r_*S*^∘^ estimation is 0.018 kJ/K/mol (Figure 4B).

We also examined how well Δ_r_*S*^∘^ correct Δ_r_*G*′^∘^ data measured at different temperatures to the same reference conditions. Specifically, we transformed Δ_r_*G*′^∘^ data to the reference state Δ_r_*G*^∘^ (298.15 K, pH 7, 0 M ionic strength, no metal ion) using equation 7. If the correction on Gibbs energies at different temperatures is beneficial, the corrected Δ_r_*G*^∘^ values should be close to the Δ_r_*G*^∘^ value at the reference conditions. For a number of case studies examined, we used Δ_r_*S*^∘^ calculated from Δ_f_ *S*^∘^ and applied the transformation to Δ_r_*G*′^∘^ data measured at different temperatures. We found a substantial decrease in Δ_r_*G*^∘^ variation with respect to temperature after applying the correction in the majority of cases (Figure 4E, 4F, S3 with yellow background). For cases where the variation in Δ_r_*G*^∘^ remains large, we calculated Δ_r_*S*^∘^ directly from the slope of Δ_r_*G*′^∘^ over T and used the value for future corrections (Table S10). The statistics of these case studies can be found in Table S13.

We also examined reactions whose Δ_r_*G*^∘^ values are sensitive to change in temperature (large Δ_r_*S*^∘^/Δ_r_*G*^∘^ ratio). A number of interesting cases in central metabolism were identified, including malate dehydrogenase, amino acid transaminase and transketolase (Table S15).

### Estimation of standard Gibbs free energy of reaction

Utilizing the curated and estimated datasets mentioned above, as well as the estimation of Δ_r_*S*^∘^, we adapted the most recent group contribution-based method, termed component contribution (10), to calculate reaction equilibrium constants for a set of 617 unique reactions in NIST TECRdb. Besides the addition of transformation of Δ_r_*G*′^∘^ across temperature, we also included 17 novel group definitions to account for compounds with new functional groups not covered by the previous component contribution method. The novel group definitions can be found in Table S11. On top of the new functionalities, we also added additional Δ_r_*G*^∘^ values of 185 reactions and Δ_f_ *G*^∘^ values of 178 compounds compared to the previous method.

We evaluated the accuracy of the updated component contribution method using repeated 10-fold cross-validation (Materials and Methods). With the additional group definitions and capability to transform Gibbs energy of reaction across temperature, we obtained a modest improvement over the latest method in median absolute residual, from 6.12 kJ/mol to kJ/mol, for a set of 432 overlapping reactions (Figure S4A) (10). Additionally, we compared the two methods by predicting Δ_r_*G*^∘^ for 185 new reactions collected in this work, using 432 reactions as training data. We found the median absolute residual from the current method (8.17 kJ/mol) is notably smaller than that from the previous work (11.47 kJ/mol) (Figure S4B). Overall, our method led to improved performance compared to the most recent group contribution method, while adding the capability to correct equilibrium constants with respect to temperature and substantially expanding the scope of predictions and thermodynamic datasets used in estimation.

## Discussion

In this work, we expanded the scope of thermodynamic calculations to more compounds and reactions with both curated and estimated data, as well as extending the group contribution methods for estimating reaction equilibrium constants to temperature conditions. We first collected and curated various types of thermodynamic data from a number of databases, including *K*′, Δ_r_*H*′^∘^, Δ_f_ *G*^∘^,Δ_f_ *H*^∘^,Δ_f_ *S*^∘^and various ion binding constants. We then estimated magnesium binding constants for 618 compounds by defining magnesium binding groups based on existing binding data. We also established a procedure to optimize magnesium binding constants using *K*′ data measured at different magnesium concentrations. Next, we applied existing thermodynamic theory with simplifying assumptions to enable the calculation of Gibbs free energy of reaction across temperature and estimated the necessary parameters (Δ_f_ *S*^∘^, Δ_r_*S*^∘^) using linear regression models. With new capabilities and new data, we utilized the updated group contribution method to calculate equilibrium constants with improve accuracy over previous work.

The curation of NIST TECRdb revealed that necessary media conditions, which influence the ionic strength and metal ion concentration corrections, were often lacking. Surprisingly, curating the literature and filling in media conditions made the resulting fit on the estimation of equilibrium constants slightly worse. This may be due to the fact that we utilized a relatively simple model to account for the effect of ionic strength on activity coefficients of aqueous electrolytes. Compared to the Debye-Hückel theory used in previous work, the Davies equation we applied in this work can handle activity coefficients of aqueous solutions at relatively high concentrations (23). However, the Davies equation still fails to account for interactions between various ions present in solution and is unable to calculate activity coefficients at temperatures other than 298.15 K. Equations with a more comprehensive handling of these thermodynamic theories are established (15–18, 35, 36), but require substantially more data than is currently available for the vast majority of compounds.

We demonstrated that magnesium binding groups identified from known magnesium binding compounds are useful features to estimate magnesium binding constants with good accuracy. However, we found a number of compounds that complex with magnesium do not contain the binding groups we defined. These compounds include nucleobases, ribonucleosides, deoxyribonucleosides, purine derivatives and small chemicals such as ammonia, thiocyanate and urea. Thus, we are unable to predict the binding constants for new compounds that fall into these categories. The approach of estimating metal binding constants by defining metal binding groups can also be applied to other metals. However, we did not perform such predictions here due to the scarcity of binding data available for other metals.

We found the overall errors in estimating Δ_r_*G*^∘^ increase with the incorporation of metal correction using curated and predicted metal binding data. We identify uncertainty in estimation of magnesium binding constants and missing binding data for other metals as primary sources of errors. Additionally, most measurements only reported total metal ion concentrations, while the metal correction formulation uses free metal ion concentrations. Therefore, additional effort is necessary to calculate free metal ion concentrations from measured data. Due to the lack of binding data and uncertainty in estimated data, an iterative approach might be taken where free metal ion concentrations calculated using the current binding data are applied to optimize the binding data, which are then fed into calculation of free metal ion concentrations. However, we still recommend the optimized magnesium binding constants to be used in metal correction, since they were obtained from well-defined cases and have proved to be effective in correcting *K*′values measured under different magnesium concentrations.

The geochemistry field has developed a complicated system to handle thermodynamic variables as a function of temperature, for a wide variety of compounds in aqueous solutions (15–21). However, the available literature only covers less than half of the compounds in NIST TECRdb. Additionally, the parameters used to calculate thermodynamic transformation across temperature are specific for different aqueous species. Therefore, we needed to estimate a large number of parameters for temperature dependent thermodynamic calculations. Due to the lack of data in necessary depth and resolution, the parameter estimation procedure resulted in large errors. Thus, with assumptions of constant enthalpy and entropy, we formulated a simplified approach to calculate temperature transformation of Gibbs energy of reaction and reduced the number of parameters needed for estimation drastically. With the incorporation of temperature transformation capabilities, we obtained similar errors in estimating Δ_r_*G*^∘^ compared to the previous method (10). Such similar errors seem to be largely due to the fact that most of the data were measured not far from 298.15 K (83.5% of the data were measured under 295.15 K to 313.15 K), resulting in minor change in correction of *K*′ to the reference conditions. However, we do observe large charge in Gibbs energy of many reactions at high temperatures (approaching 373 K), which thus may be significant for high-interest thermophile organisms such as those living in hot springs and hydrothermal vents.

Using a regression model, we predicted Δ_f_ *S*^∘^ of a comprehensive collection of compounds with high accuracy, by identifying key chemical properties such as number of atoms and partial charge. The linear correlation of other thermodynamic properties (e.g. standard molar entropy, standard partial molal volume, Δ_f_ *G*^∘^) with number of atoms has been demonstrated in previous work (37–40), but was only for compounds in the same homologous series. We found the partial charge of atoms to be useful to distinguish Δ_f_ *S*^∘^ from different homologous series, possibly due to the fact that the partial charge of atoms of the aqueous species influence its interaction with surrounding water molecules. The regression model was unable to clearly differentiate Δ_f_ *S*^∘^ of compounds within certain categories, such as monosaccharides and disaccharides. For example, the differences in Δ_f_ *S*^∘^ for fructose, mannose and sorbose are around 10 to 20 J/K/mol, while the model only predicts up to 5 J/K/mol difference, due to their similar chemical properties. Such error is not evident when evaluating the accuracy of Δ_f_ *S*^∘^ estimation, as Δ_f_ *S*^∘^ of monosaccharides are around 1000 J/K/mol. However, when calculating Δ_r_*S*^∘^ of the isomerization reaction between monosaccharides, we found that the errors of Δ_f_ *S*^∘^ prediction and the calculated Δ_r_*S*^∘^ values are around the same order of magnitude. We observed this issue to be prevalent for a number of reactions in NIST TECRdb. Thus, we developed another regression model to estimate Δ_r_*S*^∘^ values from existing data, rather than calculating from Δ_f_ *S*^∘^ predictions.

Taken together, we expanded the framework to calculate Gibbs free energy of reaction under different conditions, with curated and estimated data enabling calculations across different temperatures and metal concentrations. This effort expands opportunities toward an understanding of thermodynamic factors underlying metabolic network and function in biological systems. This area has generated a number of exciting results, such as the discovery that amino acid biosynthesis, which is endergonic at surface conditions, is exergonic under the conditions of life in hydrothermal vents (41). Another recent effort proposed proteomic constraints due to thermodynamic bottlenecks as a critical factor underlying use of the Entner-Doudoroff pathway (4). As methods for estimating the thermodynamic properties of metabolic networks continue to improve, these efforts are likely to be increasingly fruitful in uncovering the physical constraints driving the evolution of metabolic networks.

## Conclusion

The work here provides an updated group contribution method with an expanded set of thermodynamic data and extended capabilities to calculate equilibrium constants as a function of temperature. We collected and curated thermodynamic data for compounds and reactions from a number of databases and primary literature sources. We defined magnesium binding groups from existing data and estimated magnesium binding constants for 618 compounds through a linear regression model. We also established a simple yet well-justified framework, which included formulations derived from existing theory and the necessary parameters (Δ_f_ *S*^∘^, Δ_r_*S*^∘^), to calculate equilibrium constants as a function of temperature. Taken together, this work fills a gap in previous group contribution methods to calculate equilibrium constants to temperature conditions and better correct for metal ion binding. These efforts should facilitate the growing number of applications to apply thermodynamic principles to better understand cell metabolism.

## Author Contributions

BD and DCZ conceived and designed the study. BD, ZZ, SG, JTY and DCZ collected the data. BD and DCZ performed the analysis. BD, ZZ, SG, JTY, BOP and DCZ wrote the paper. BD and JTY wrote the Supporting Material. All authors read and approved the final content.

## Acknowledgments

We would like to thank Nikolaus Sonnenschein for valuable discussions. This work was supported by the Novo Nordisk Foundation Grant Number NNF10CC1016517.

## Supporting Citations

References (6, 9–11, 17, 19, 23, 28, 42–53) appear in the Supporting Material and Supporting Data.

