## Supplementary Materials for "Temperature-dependent estimation of Gibbs energies using an updated group contribution method"

BD, ZZ, SG, JTY, BOP, DCZ

### Transformation of standard Gibbs free energy of formation ($\Delta_\mathrm{f}G^\circ$) of aqueous species across temperature

Considering constant pressure, the standard Gibbs free energy of formation of an aqueous species at a given temperature T and the reference temperature $T_\mathrm{r}$ (298.15 K) can be written as $\Delta_\mathrm{f}G^\circ_T$ and $\Delta_\mathrm{f}G^\circ_{T_\mathrm{r}}$, respectively. Based on the Second law of thermodynamics,

$$\Delta_\mathrm{f}G^\circ_T = \Delta_\mathrm{f}H^\circ_T - T\Delta_\mathrm{f}S^\circ_T \quad [1]$$

$$\Delta_\mathrm{f}G^\circ_{T_\mathrm{r}} = \Delta_\mathrm{f}H^\circ_{T_\mathrm{r}} - T_\mathrm{r}\Delta_\mathrm{f}S^\circ_{T_\mathrm{r}}. \quad [2]$$

Subtracting Equation 1 by Equation 2, we get

$$\begin{aligned}\Delta_\mathrm{f}G^\circ_T = \Delta_\mathrm{f}G^\circ_{T_\mathrm{r}} + (\Delta_\mathrm{f}H^\circ_T - \Delta_\mathrm{f}H^\circ_{T_\mathrm{r}}) - \\ T(\Delta_\mathrm{f}S^\circ_T - \Delta_\mathrm{f}S^\circ_{T_\mathrm{r}}) - (T - T_\mathrm{r})\Delta_\mathrm{f}S^\circ_{T_\mathrm{r}}\end{aligned} \quad [3]$$

Using the definition of enthalpy and entropy in terms of heat capacity at constant pressure (1, 2), Equation 3 is expressed as

$$\begin{aligned}\Delta_\mathrm{f}G^\circ_T = \Delta_\mathrm{f}G^\circ_{T_\mathrm{r}} + \int_{T_\mathrm{r}}^{T} C_{P_\mathrm{r}}dT - \\ T\int_{T_\mathrm{r}}^{T} C_{P_\mathrm{r}}d\ln T - (T - T_\mathrm{r})\Delta_\mathrm{f}S^\circ_{T_\mathrm{r}}\end{aligned} \quad [4]$$

where $C_{P_\mathrm{r}}$ is the heat capacity of the aqueous species at $T_\mathrm{r}$. It is worth mentioning that the formulation above is slightly different from that in the geochemistry literature (1, 2), where we replaced $S^\circ_{T_\mathrm{r}}$ with $\Delta_\mathrm{f}S^\circ_{T_\mathrm{r}}$.

Based on Shock et al. (2), the heat capacity of an aqueous species is a function of temperature and depends on three parameters $c_1$, $c_2$, and $\omega$, which are different for different aqueous species. We found that heat capacities of aqueous species at different temperatures generally vary in a small range from their values at $T_\mathrm{r}$. Specifically, we examined a total of 399 compounds with data available (3) and found that their $C_P$ values at different temperatures vary maximally around 16% from their $C_{P_\mathrm{r}}$ values, for the temperature range we are working with (283.5 K to 360.5 K). The temperature range is based on temperatures of measured data in TECRdb (4). Additionally, the maximum variation in $C_P$ across temperature is smaller than that across different compounds, as shown in Figure S1A. Thus, given the assumption that heat capacity is a constant with respect to temperature, we take integrals in Equation 4 and get

$$\begin{aligned}\Delta_\mathrm{f}G^\circ_T = \Delta_\mathrm{f}G^\circ_{T_\mathrm{r}} + C_{P_\mathrm{r}}(T - T_\mathrm{r}) - \\ TC_{P_\mathrm{r}}\ln\left(\frac{T}{T_\mathrm{r}}\right) - (T - T_\mathrm{r})\Delta_\mathrm{f}S^\circ_{T_\mathrm{r}}.\end{aligned} \quad [5]$$

Combining the terms involving $C_{P_\mathrm{r}}$, we have

$$\begin{aligned}\Delta_\mathrm{f}G^\circ_T = \Delta_\mathrm{f}G^\circ_{T_\mathrm{r}} + \left[T - T_\mathrm{r} - T\ln\left(\frac{T}{T_\mathrm{r}}\right)\right]C_{P_\mathrm{r}} - \\ (T - T_\mathrm{r})\Delta_\mathrm{f}S^\circ_{T_\mathrm{r}}\end{aligned} \quad [6]$$

From Equation 6, we have the term involving $C_{P_\mathrm{r}}$ and the term involving $\Delta_\mathrm{f}S^\circ_{T_\mathrm{r}}$ together affecting the change of standard Gibbs free energy of formation across temperature.

Comparing the coefficients in front of $C_{P_\mathrm{r}}$ and $\Delta_\mathrm{f}S^\circ_{T_\mathrm{r}}$ at different temperatures, we found that the $C_{P_\mathrm{r}}$ coefficient is much smaller than that of $\Delta_\mathrm{f}S^\circ_{T_\mathrm{r}}$ (Figure S1B). The ratio of $C_{P_\mathrm{r}}$ coefficient to $\Delta_\mathrm{f}S^\circ_{T_\mathrm{r}}$ coefficient is at most 0.025 for the most frequent temperatures of TECRdb measured data (295.5 K to 313.5 K), and at most 0.1 in the overall temperature range of interest.

Given that $C_{P_\mathrm{r}}$ and $\Delta_\mathrm{f}S^\circ_{T_\mathrm{r}}$ of the same aqueous species are generally on the same order of magnitude (Figure S1C) and $C_{P_\mathrm{r}}$ coefficient is much smaller than that of $\Delta_\mathrm{f}S^\circ_{T_\mathrm{r}}$, it is reasonable to neglect the term involving $C_{P_\mathrm{r}}$ in Equation 6. Thus, we have

$$\Delta_\mathrm{f}G^\circ_T = \Delta_\mathrm{f}G^\circ_{T_\mathrm{r}} - (T - T_\mathrm{r})\Delta_\mathrm{f}S^\circ_{T_\mathrm{r}} \quad [7]$$

to transform the standard Gibbs free energy of formation of an aqueous species across temperature.

### Equilibrium constant as a function of pH, temperature, ionic strength and metal ion concentration

In aqueous solutions, each compound exists as several different pseudoisomer forms distributed according to the Boltzmann distribution. The pseudoisomer forms refer to the different protonation and ion bound states of the same compound (5, 6). For example, the pseudoisomer forms of orthophosphate include but are not limited to $PO_4^{3-}$ and $MgPO_4^-$. Adapted from Alberty (6) and the formulation in the last section, the standard transformed Gibbs free energy of formation of pseudoisomer $i$ ($\Delta_\mathrm{f}G'^\circ_i$) of a given compound under certain pH, temperature ($T$), ionic strength ($I$) and metal ion concentration (pM) is expressed as

$$\begin{aligned}\Delta_f G_i'^\circ = \Delta_f G_i^\circ(I=0,T_r) - (T-T_r)\Delta_f S_i^\circ + \\ N_H(i)RT\ln(10)\text{pH} - N_M(i)(\Delta_f G_M^\circ(T) - \\ RT\ln(10)\text{pM}) - RT\alpha(z_i^2 - N_H(i))\left(\frac{\sqrt{I}}{1+\sqrt{I}} - 0.3I\right)\end{aligned} \qquad [8]$$

where $\Delta_f S_i^\circ$ is the standard entropy change of formation of pseudoisomer $i$ at 298.15 K, $z_i$, $N_H(i)$, and $N_M(i)$ are the charge, number of hydrogen atoms and number of metal ions M bound to pseudoisomer $i$ (due to availability of metal binding data, we only handle pseudoisomer form bound with at most one type of metal ion), $\Delta_f G_M^\circ(T)$ is the standard Gibbs free energy of formation of aqueous ionic metal species M at $T$ (can be calculated using equations and data from Shock et al. (2)), pM (pM $= -\log_{10}[M^{m+}]$) is the potential of ionic metal species M with concentration [M] and charge +m in aqueous solutions, and $\alpha$ is the Debye-Hückel Constant and is temperature dependent (6). The correction on ionic strength is based on Davies equation, which is an empirical extension of Debye–Hückel theory and can be used to calculate activity coefficients of electrolytes at relatively high ion concentrations (7).

The standard transformed Gibbs free energy of formation of the compound ($\Delta_f G_j'^\circ$) can be calculated based on the energies of its pseudoisomer forms using Legendre transform (6):

$$\Delta_f G_j'^\circ = -RT\ln\left\{\sum_{i=1}^{N_{\text{iso}}}\exp\left[-\frac{\Delta_f G_i'^\circ}{RT}\right]\right\}. \qquad [9]$$

The standard transformed Gibbs free energy of reaction ($\Delta_r G'^\circ$) can thus be calculated based on the energies of its participating compounds ($\Delta_f G_j'^\circ$) and their corresponding stoichiometries ($r_j$) in the reaction

$$\Delta_r G'^\circ = \sum_{j=1}^{N} r_j \Delta_f G_j'^\circ \qquad [10]$$

Thus, we are able to calculate thermodynamics of the reaction as a function of pH, temperature, ionic strength and metal ion concentrations.

However, when we calculated the standard entropy change of reaction ($\Delta_r S^\circ$) from $\Delta_f S^\circ$ of the participating species, we found that $\Delta_f S^\circ$ values are usually larger in order of magnitude compared to $\Delta_r S^\circ$ values (Figure S1D). The uncertainties in $\Delta_r S^\circ$ calculations based on those in $\Delta_f S^\circ$ estimations (main text, Figure 3) are usually around the same order of magnitude as $\Delta_r S^\circ$ values themselves. Thus, the transformation of Gibbs energy of reaction across temperature using $\Delta_f S^\circ$ estimations is likely to be error-prone. To address this issue, we estimated $\Delta_r S^\circ$ at $T_r$ (298.15 K) from existing data (main text, Figure 4), rather than calculating from $\Delta_f S^\circ$ estimations. We also modified the formulation of calculating Gibbs energy of reaction as a function of temperature, using $\Delta_r S^\circ$ at $T_r$

$$\Delta_r G_T'^\circ = \Delta_r G_{T_r}'^\circ - (T-T_r)\Delta_r S_{T_r}^\circ \qquad [11]$$

The reaction equilibrium constant can thus be calculated through the equation

$$\Delta_r G'^\circ = -RT\ln K'. \qquad [12]$$

The above procedures can also be used to transform the measured equilibrium constants to $\Delta_r G^\circ$ at the reference state (298.15 K, pH 7, 0M ionic strength, no metal ion), by applying corrections on pH, ionic strength and metal ion concentrations as in Equation 8 and correction on temperature as in Equation 11. Then, we can use corrected $\Delta_r G^\circ$ data to estimate $\Delta_r G^\circ$ and $\Delta_f G^\circ$ for new reactions and compounds, based on the latest group contribution method, termed component contribution (8).

### Example of binding constant and binding polynomial formulation

We introduce the concept of binding constant and describe its relationship with the binding polynomial. Binding constant describes the equilibrium of binding and unbinding reaction between a receptor (compound) and a ligand (proton, metal ion). Here, we specifically refer the binding constant to be the equilibrium constant of the unbinding step. For example, a reactant is composed of three ion bound states: A (with least hydrogens and metal ions bound), HA (A bound with $H^+$), MgHA (A bound with $H^+$ and $Mg^{2+}$). There are two binding steps between A and MgHA: HA $\rightleftharpoons$ A + $H^+$ and MgHA $\rightleftharpoons$ HA + $Mg^{2+}$. The respective binding constants are

$$K_1 = \frac{[A][H^+]}{[HA]} \qquad [13]$$

$$K_2 = \frac{[HA][Mg^{2+}]}{[MgHA]} \qquad [14]$$

For practical purposes, it is more convenient to express the logarithmic form of the constants, where $pK_1 = -\log_{10}K_1$ and $pK_2 = -\log_{10}K_2$. Based on the type of ligand, $pK_1$ is known as the acid dissociation constant ($pK_a$) and $pK_2$ is the stability constant for magnesium binding ($pK_{Mg}$). The logarithmic form of the binding constant is what we used for estimation in regression models and calculation in the group contribution framework.

Binding polynomial gives the partition of a reactant between various aqueous species that make it up. Binding polynomial is the measure of the difference in Gibbs energy between one ion bound state and another. For convenience of calculation, we usually write the binding polynomial of an ion bound state with respect to the one with the least hydrogens and metal ions bound. Thus, the binding polynomial $P$ of MgHA is defined as (6):

$$P = \frac{[A] + [HA] + [MgHA]}{[A]} \qquad [15]$$

Substituting Equation 13 and 14 into 15, we get

$$P = 1 + \frac{[H^+]}{K_1} + \frac{[H^+][Mg^{2+}]}{K_1K_2} \qquad [16]$$

The energy difference between A and MgHA is $-RT\ln P$, which can be used in Equation 6 of the main text and calculate $\Delta_f G'^\circ$ of the reactant. Therefore, binding polynomial can be expressed in terms of proton and metal ion concentrations, as well as the binding constants of different binding steps. This example can be extended to any other ion bound states with defined number of hydrogens and metal ions bound.

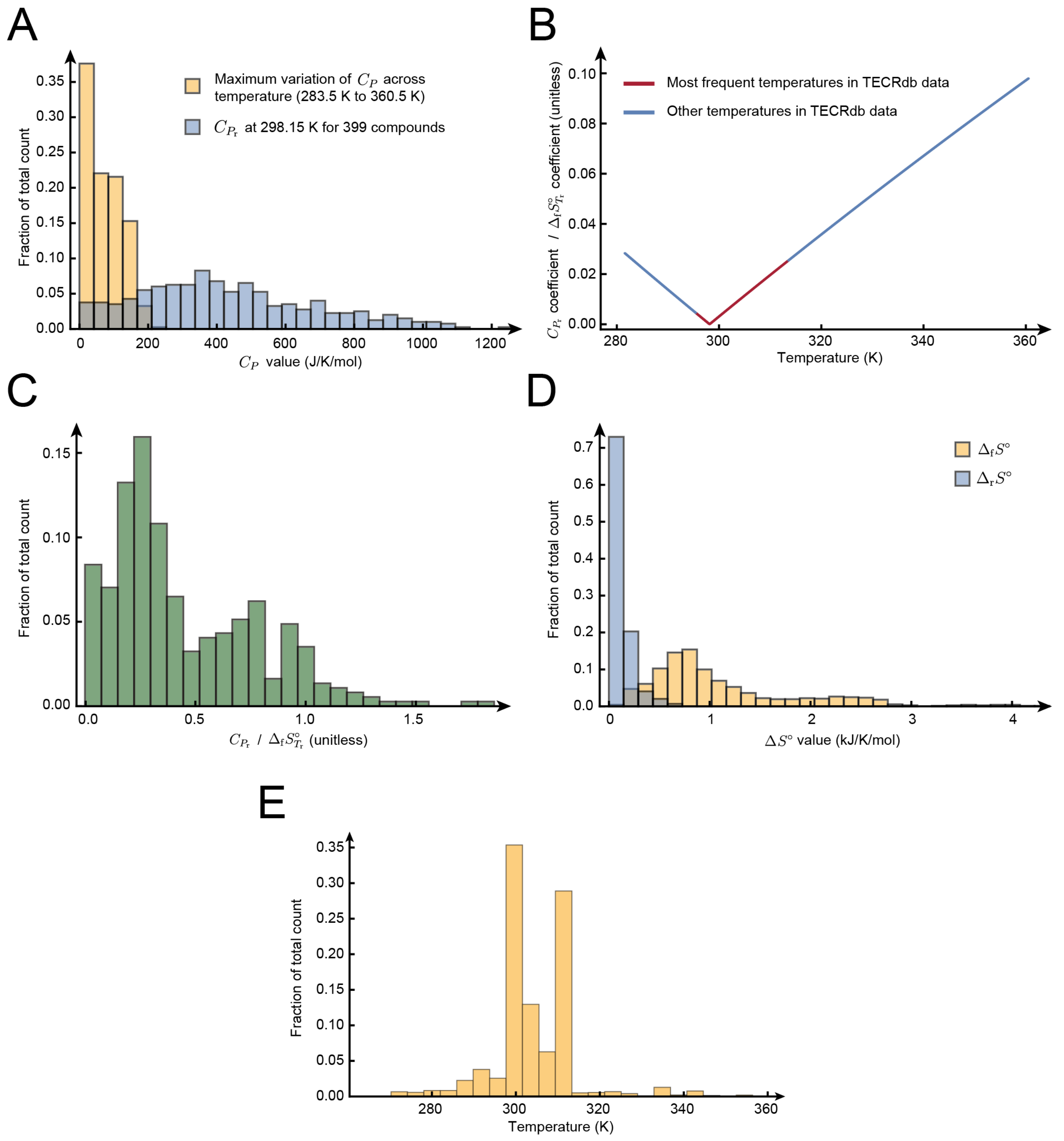


**Fig. S1.** (A) Order of magnitude comparison between maximum variation of $C_P$ across temperature vs. $C_{P_\mathrm{r}}$ at 298.15 K for 399 compounds (3). (B) The ratio of $C_{P_\mathrm{r}}$ coefficient to $\Delta_\mathrm{f} S^\circ_{T_\mathrm{r}}$ coefficient as a function of temperature. The temperature range is based on the distribution of temperatures for measured equilibrium constants in TECRdb. (C) The distribution of $C_{P_\mathrm{r}}/\Delta_\mathrm{f} S^\circ_{T_\mathrm{r}}$ of the same aqueous species from a total 370 aqueous species collected. (D) Order of magnitude comparison between $\Delta_\mathrm{f} S^\circ$ and $\Delta_\mathrm{r} S^\circ$. We collected a total of 669 $\Delta_\mathrm{f} S^\circ$ values and 148 $\Delta_\mathrm{r} S^\circ$ values for comparison. (E) Distribution of temperature for all measured equilibrium constants in TECRdb.

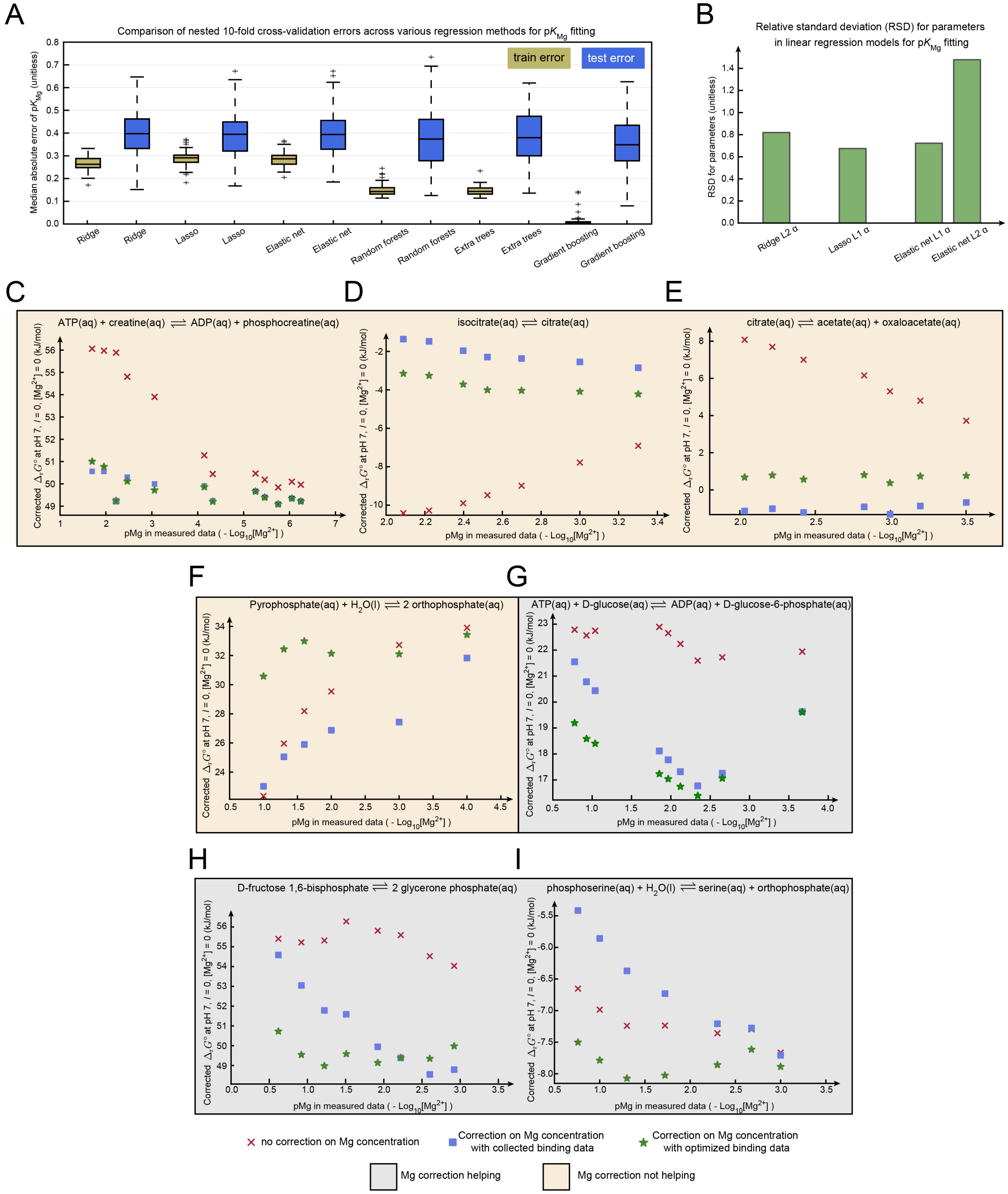


**Fig. S2.** (A) Training and testing errors of nested 10-fold cross validation on magnesium (Mg) binding data using the ridge regression, lasso regression, elastic net regularization, random forests, extra trees and gradient boosting. We repeated cross-validation 5 times by splitting all Mg binding data into different subdivisions. We included a total of 140 Mg binding data points and 128 features including metal binding groups, the partial charge, and molecular properties from ChemAxon and RDKit. (B) Relative standard deviation (RSD) for parameters in linear regression models used for Mg binding fitting. We calculated the mean and standard deviation of parameters selected by the inner loops of nested cross validation (repeated 5 different times). We used RSD (standard deviation/mean) to assess the relative variability of the parameters and model stability in Mg binding fitting. (C-I) Case studies on corrected $\Delta_r G^\circ$ values of different reactions calculated from equilibrium constants ($K'$) measured at different Mg concentrations. We applied different corrections to transform $\Delta_r G'^\circ$ (calculated from $K'$) to $\Delta_r G^\circ$ values: no correction on varying Mg concentrations (red Xs), correction on varying Mg concentrations using collected binding constants (blue squares), correction on varying Mg concentrations using optimized binding constants (green stars). The reaction whose $\Delta_r G'^\circ$ data are used to optimize the binding constants can be found in Table S12. Panels with yellow background are cases where applying original Mg binding constants reduces the variation in $\Delta_r G^\circ$, while those with grey background are cases where using original Mg binding data did not help. $\mathrm{p}K_{\mathrm{Mg}}$: magnesium binding constant.

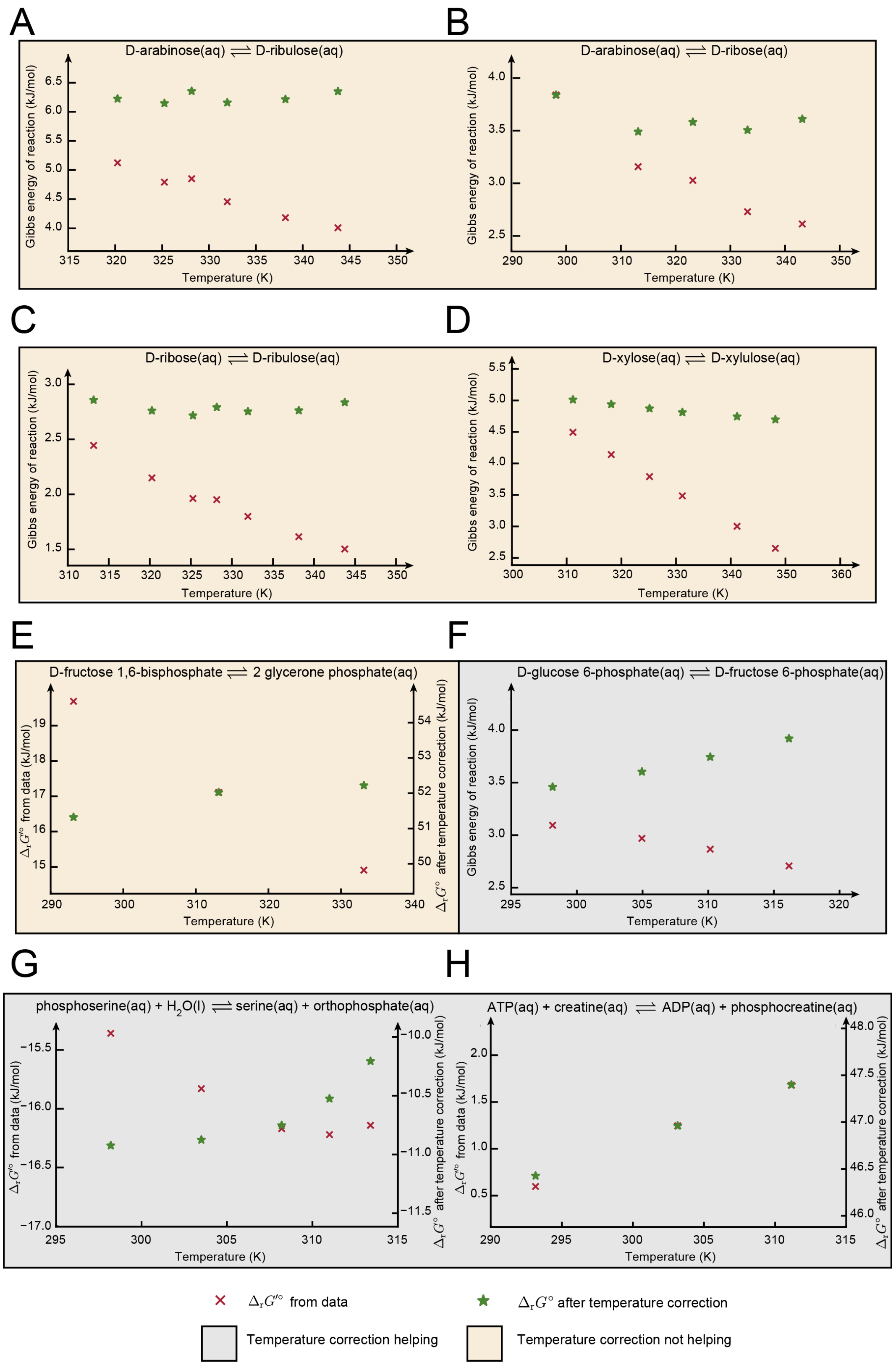


**Fig. S3.** Case studies on corrected $\Delta_r G^\circ$ (green stars) vs. $\Delta_r G'^\circ$ (red Xs) values (calculated from $K'$) of different reactions measured at different temperatures. We calculated $\Delta_r G'^\circ$ from $K'$ and compared with $\Delta_r G^\circ$ values after applying temperature correction. Panels with yellow background are cases where applying temperature correction reduces the variation in $\Delta_r G^\circ$, while those with grey background are cases where temperature correction did not help.

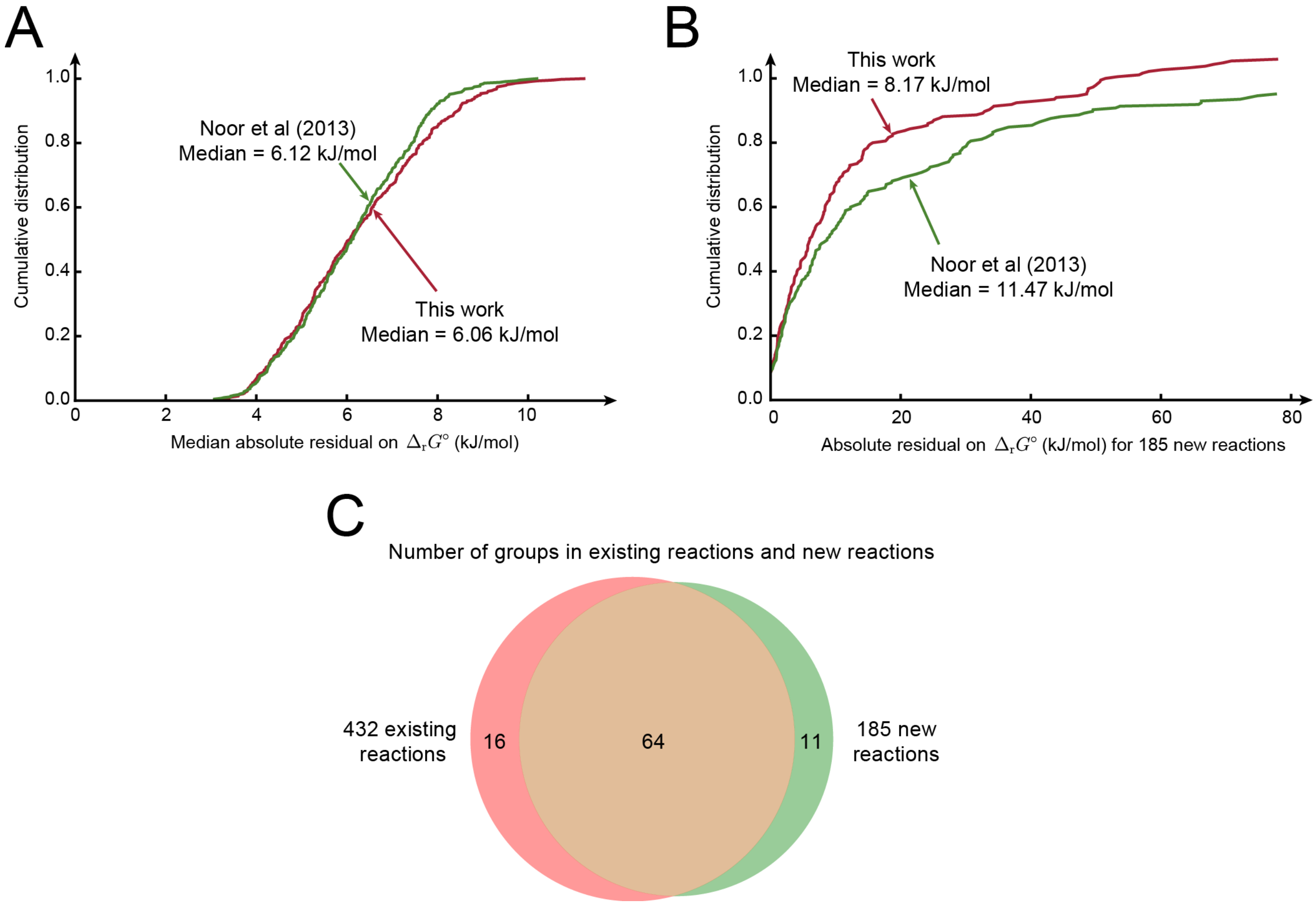


**Fig. S4.** (A) Comparison of absolute residuals on estimating $\Delta_r G^\circ$ for 432 overlapping reactions between the previous group contribution method (8) and the current method. We performed 100 repetitions of 10-fold cross-validation using the previous method and the current method. (B) Comparison of absolute residuals on estimating 185 new reactions in the current method. We calculated $\Delta_r G^\circ$ for 185 new reactions by constructing the group contribution model using $\Delta_r G^\circ$ values of 432 overlapping reactions from the previous method and the current method. We then calculated the absolute residual between estimated $\Delta_r G^\circ$ and $\Delta_r G^\circ$ data for those 185 reactions. (C) Comparison of group coverage between 432 reactions in the previous group contribution method and 185 new reactions added in the current method.
